# Activation of ARTD2/PARP2 by DNA damage induces conformational changes relieving enzyme autoinhibition

**DOI:** 10.1101/2020.11.24.395475

**Authors:** Ezeogo Obaji, Mirko M. Maksimainen, Albert Galera-Prat, Lari Lehtiö

**Author notes:** Corresponding author: Lari Lehtiö.

## Abstract

Human ARTD2/PARP2 is an ADP-ribosyltransferase which, when activated by 5’- phosphorylated DNA ends, catalyzes poly-ADP-ribosylation of itself, other proteins and DNA. A crystal structure of ARTD2 in complex with an activating 5’-phosphorylated DNA shows that the WGR domain bridges the dsDNA gap and joins the DNA ends. This DNA binding results in major conformational changes, reorganization of helical fragments, in the ARTD2 regulatory domain. Comparison of ARTD1-3 crystal structures reveal how binding to a DNA damage site leads to formation of a catalytically competent conformation capable of binding substrate NAD^+^ and histone PARylation factor 1 changing the ARTD2 residue specificity from glutamate to serine when initiating DNA repair processes. The structure also reveals how the conformational changes in the autoinhibitory regulatory domain would promote the flexibility needed by the enzyme to reach the target macromolecule for ADP-ribosylation.

## Main

DNA damage is a common event in cells; about 10^4^-10^5^ DNA lesions per day are detected by enzymes of the human ADP-ribosyltransferase family (ARTD1-3/PARP1-3). In the nucleus, the main enzymes carrying out poly-ADP-ribosylation are ARTD1 and ARTD2. These enzymes bind to the damaged DNA and subsequently generate poly-ADP-ribose chains (PAR) that act as recruitment signals for a range of DNA repair factors^1–3^. The role of ADP-ribosylation is established in single strand break repair (SSBR) and in alternative non-homologous end joining (aNHEJ) mechanisms, where the key proteins involved in the repair pathways are known to be recruited to the site of DNA damage in PAR dependent manner^4–9^. Poly-ADP-ribosylation initiates also chromatin remodeling through PAR binding ALC1 (amplified in liver cancer 1)^10,11^.

ARTD1-3 enzymes have similar domain organization at the C-terminal catalytic part consisting of an ADP-ribosyltransferase domain, regulatory domain (RD) and a WGR domain shown to participate in DNA binding. They however differ in their N-terminal parts as ARTD1 contains a BRCA1 C Terminus domain (BRCT) and three zinc-fingers (ZnFs), whereas the N-termini of ARTD2 and 3 are intrinsically disordered^12,13^. It has been shown that the WGR domain of ARTD2 is a key to the DNA damage recognition^12–14^. *In vitro,* ARTD1 can be activated by multiple forms of DNA damage mimicking oligonucleotides^15,16^, while ARTD2 and ARTD3 are specifically activated by 5’-phosphorylated DNA breaks^12,14,17^. While ARTD1-3 employ different mechanisms in the DNA damage recognition, it is thought that their activation will be similar as they all contain an autoinhibitory RD domain. RD domain, in the inactive state, covers the active site and prevents binding of the substrate NAD^+^ to the catalytic domain^16,18^.

Multiple structures of the catalytic domain of ARTDs^19–22^ and of individual domains binding to DNA are available for ARTD1 and ARTD2^14,23–26^. For ARTD1 activation, a conformational change at the catalytic fragment is necessary^16,18^ and that is triggered by the sequential reorganization of the protein domains on the detected DNA damage site^26^. Furthermore, binding of histone PARylation factor 1 (HPF1) to the ARTD domain results in a joint catalytic site with changed specificity from glutamate and aspartate to serine^27^. Despite the recently reported structures and HXMS studies, it is still poorly understood how the binding of the enzyme to an activating DNA molecule can trigger a robust up to 500-fold activity increase^5,12,14,26^. Here we elucidate this process by describing the structural basis of ARTD2 DNA dependent activation. Activation induces major conformational changes in domain structure and reordering of the secondary structure elements of the RD domain to release the enzyme from an autoinhibited state.

## Results

### Crystal structure of ARTD2-DNA

We determined an ARTD2 crystal structure consisting of the WGR domain and catalytic fragment (ARTD2WGR-RD-ART: residues 90-585) in complex with an activating double stranded DNA mimicking a damaged oligonucleotide. The structure was obtained by using a multicrystal approach where 27 small datasets were merged in order to achieve a complete data at 2.8 Å resolution (**Table 1**). The asymmetric unit contains one ARTD2_WGR-RD-ART_ molecule with one dsDNA (DNA-1, **Table S1**), while the biological unit has 2:2 stoichiometry in solution (**Fig. S1**). The complex is formed by two dsDNA oligonucleotides joined together at the phosphorylated ends by two flanking proteins (**Fig. 1A**). The WGR domain of ARTD2 interacts with the 5’-phosphorylated site and with DNA on both sides of the nick as observed with the isolated WGR domain^14^. The two DNA molecules and two ARTD2s are related through a 2-fold symmetry axis. As ARTD2 is robustly activated by a 5’-phosphorylated single strand break, the ARTD2 molecules represent monomeric proteins detecting a DNA nick individually.

**Table 1:**
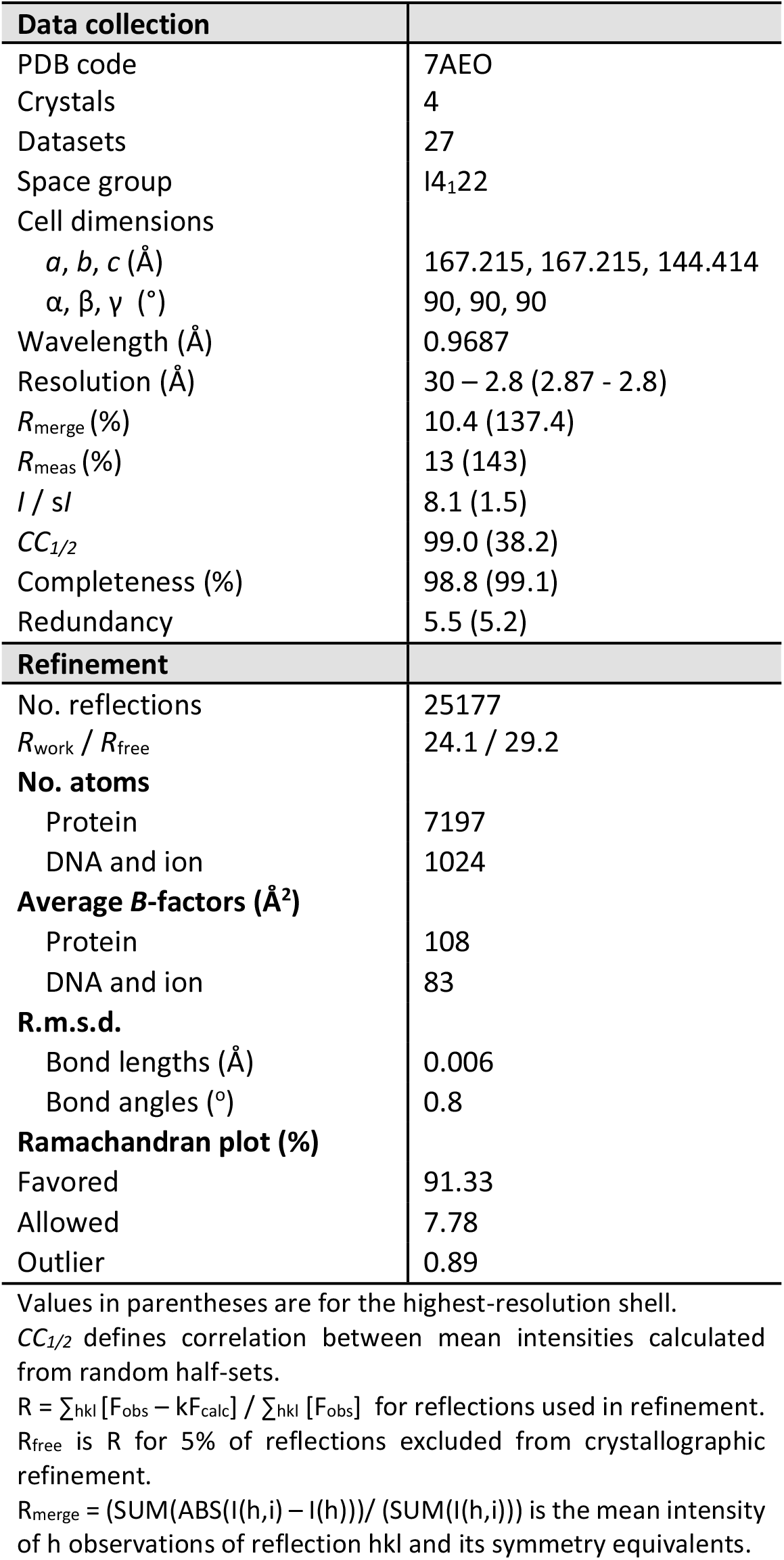
Data collection and structure refinement statistics for the ARTD2_WGRCAT_ + 5’P- dsDNA complex structure.

**Figure 1.**
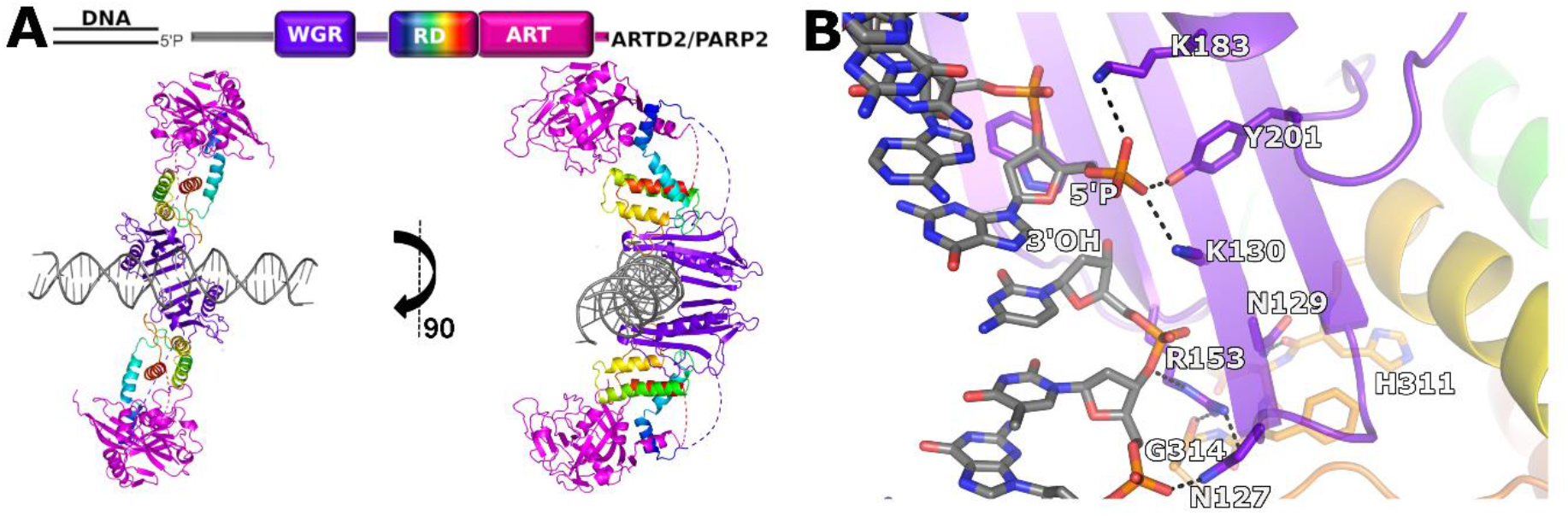
Crystal structure of ARTD2 activated by binding to DNA damage. **(A)** The biological unit as observed in solution contains two dsDNA molecules joined together by two ARTD2 molecules binding to the two nicks formed at the DNA break. The WGR domain is shown in purple, C-terminal transferase domain in magenta and RD from blue to red from N-terminus to C-terminus. **(B)** WGR domain and DNA interaction interface.

The WGR domain detects the phosphorylated DNA end coordinated by Trp151, Lys130, Lys183 and Tyr201. Tyr201 (Phe638 in ARTD1) forms a hydrogen bond with the phosphate, whereas Arg153 and Asn127 (Gly565 in ARTD1) interact with the n-1 nucleotide from the 3’end and bridge the connection to Gly314 of the RD domain **(Fig. 1B)**. The orientation of the residues is similar to the DNA complexes of the isolated WGR domain^14^ and to the recently reported Cryo-EM structure^28^.

### Opening of DNA end and interaction with ARTD2 catalytic domain

Interestingly, in addition to binding to the phosphorylated DNA break by the WGR domains, the crystal symmetry showed that the catalytic site was also in contact with the DNA, namely the un-phosphorylated end, where the last A-T base pair is opened up (**Fig. 1A**). The 3’- adenosine ribose interacts with the helix lining the active site and its ribose makes only one hydrogen bond to Asp396, while the 5’-thymine binds to the active site (**Fig. 2AB**) and locates in the nicotinamide binding site between two tyrosine side chains (**Fig. 2C**). The base forms typical hydrogen bonds with Gly429 and Ser470 like the amide group of the PARP inhibitors and of the substrate mimicking analog (**Fig. 2D**). In addition, the ribose and the phosphate of the nucleotide bind to the same regions in the active site where the substrate NAD^+^ is expected to interact. This implies that thymine could also act as an inhibitor of the enzyme. Therefore, we performed an inhibition assay using thymine, thymidine and thymidine monophosphate and the result showed an IC_50_ of 50 μM for thymine (**Fig. S2**). Thymidine has an even lower IC_50_ of 14 μM, while thymidine monophosphate is not a potent ARTD2 inhibitor with an IC_50_ of 681 μM, which indicates that most of the binding energy comes from the nicotinamide mimicking thymine and the ribose.

**Fig. 2.**
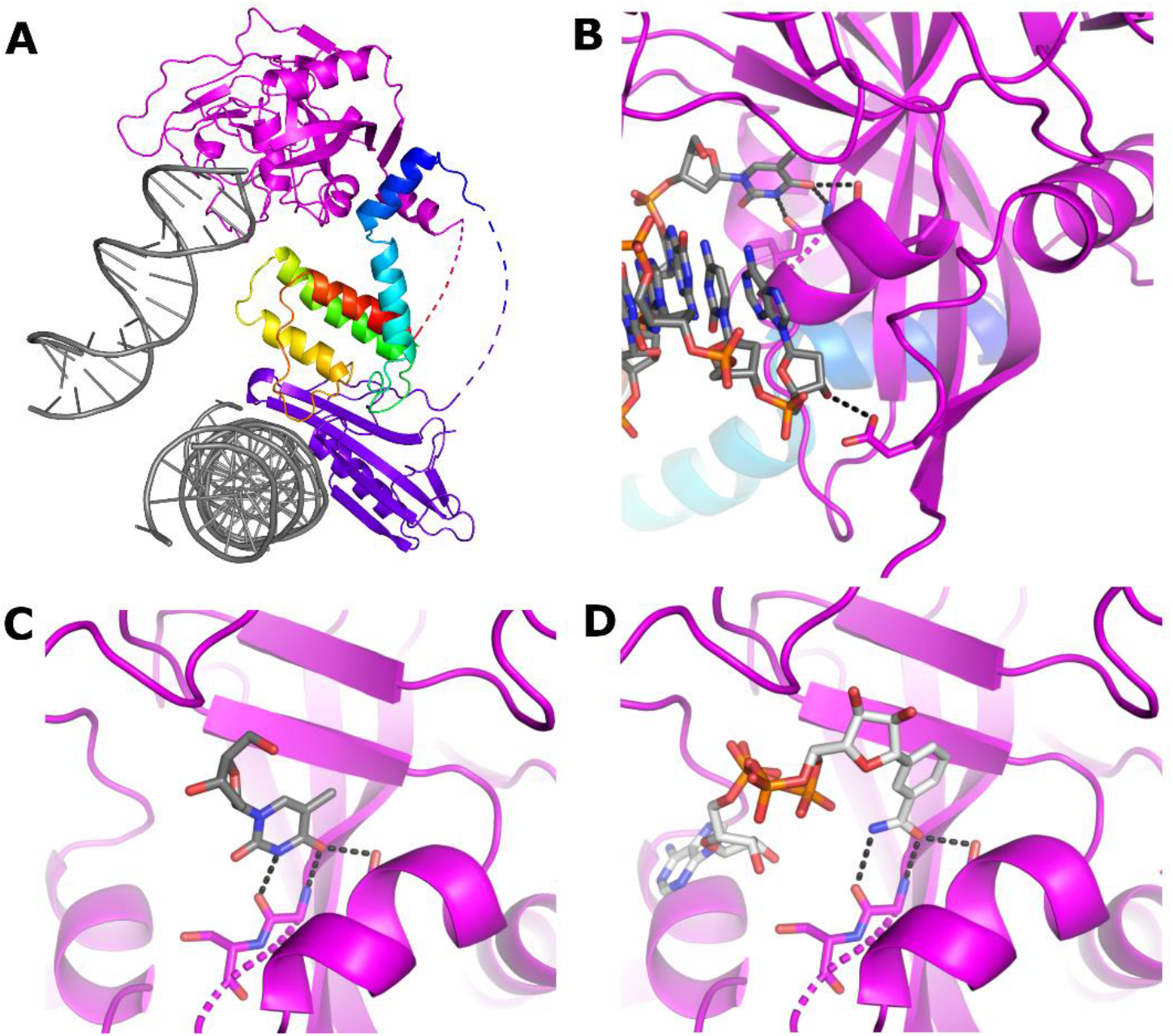
Binding of the unphosphorylated DNA end to the catalytic site. (**A**) DNA end binding to the catalytic domain and formation of a crystal contact. (**B**) Interaction of the DNA 3’and 5’-ends with the transferase domain. (**C**) Close up view of the 5’-thymidine in the nicotinamide binding site. (**D**) Binding of an NAD^+^ analog to the active site superimposed from the ARTD1 complex structure (PDB id. 6BHV).

In addition, to determine how the activation capacity of the particular DNA used in the crystallization would affect the PAR synthesis of ARTD2, we measured the DNA dependent activity of the enzyme using the same DNA as in the crystal structure as well as other model DNAs. Indeed the DNA used in the crystallization showed reduced PAR synthesis compared to nicked hairpin DNA and a longer form of dumbbell hairpin DNA (**Fig. S2**). Conversely, when we substituted the thymine in the DNA used for the crystallization with guanine and used DNA of different lengths (DNA-4 and DNA-5) PAR synthesis increased (**Fig. S2**). The above indicates that a terminal A-T pair of the DNA would also inhibit ARTD2 in solution.

### Local unfolding of ARTD2 RD domain and autoinhibitory effect

Comparison of the ARTD2WGR-RD-ART structure with individual domain structures and the ARTD1 DNA complex reveals major conformational changes that occur upon DNA binding. These include a movement of the ART domain with respect to the rest of the protein and reorganization of the RD domain. The RD reorganization leads to opening of the catalytic site and local unfolding, especially of helix α5 packing in the inactive state against the catalytic domain (**Fig. 3A-C**). This is in line with previous studies using HXMS that show local conformational changes are required to unfold the RD helix and this in turn inhibits enzymatic activity by covering the NAD^+^ binding site^15,16,18^. In addition, the crystal structure revealed that, while the RD region close to the DNA and interacting with the WGR domain remains unaffected, the conformational changes involve more than the unfolding of a helical fragment as both the N- and C-terminal helices of the RD region undergo major reorganization (**Fig. 3A-C**). Upon DNA binding, helix α5, covering the active site in the inactive conformation, is divided into two parts at Gly338, and the helices are completely reorganized. Subsequently the catalytic transferase domain moves ~11 Å away from the DNA and it is rotated and translated implying mobility in solution (**Fig. 3A-C**).

**Figure 3.**
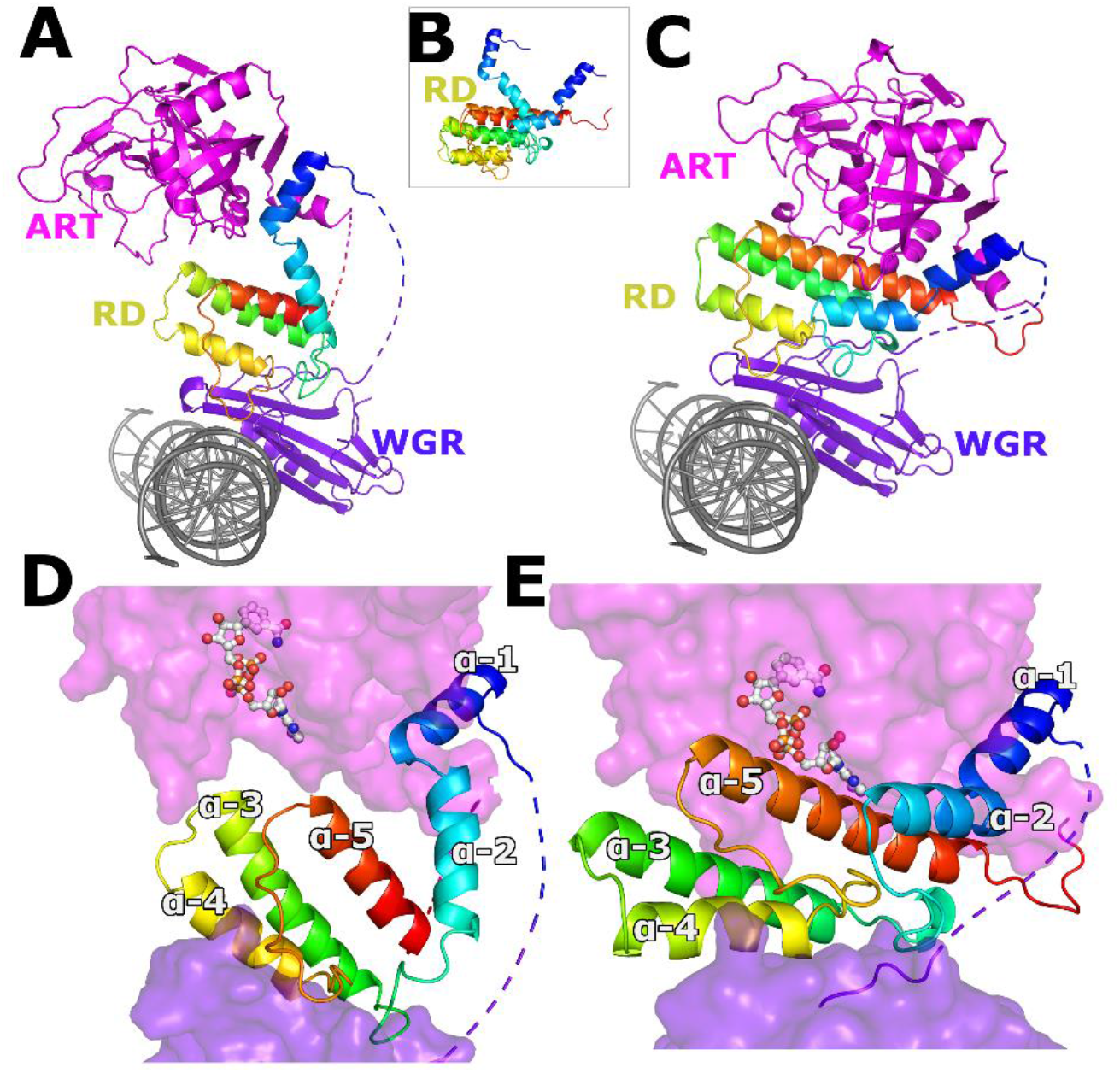
Conformation changes upon ARTD2 activation. **(A)** Activated ARTD2 structure. **(B)** Close up view comparing the RD domain in the active and inactive conformations. **(C)** A model of ARTD2 binding to DNA in an inactive conformation where the ARTD2RD-CAT crystal structure (PDB id. 4TVJ) was positioned as observed on the ARTD1 structure (PDB id. 4DQY). **(D)** Illustration showing how DNA activation releases autoinhibition of the catalytic domain allowing substrate binding (BAD shown in sticks, PDB id. 6BHV). (**E**) RD in an inactive state blocks the binding of the substrate NAD^+^.

### Mechanism of substrate binding by ARTD2 upon DNA activation

Recently, a crystal structure of a constitutively active transferase fragment consisting of only an ARTD domain was solved in complex with an unhydrolyzable substrate analog benzamide adenine dinucleotide (BAD)^18^. This ARTD1 crystal structure was used to model substrate binding in ARTD2 and compare its accessibility in the inactive and active conformations. The ARTD2 inactive model is not compatible with substrate binding (**Fig. 3E**) due to steric effects caused by RD residues (**Fig. S3**). In the active conformation, the RD helices have moved and exposed the active site of the transferase domain, which is now fully capable of binding the substrate NAD^+^ (**Fig. 3D**). ARTD2 is known to catalyze automodification and ADP-ribosylate itself in *cis* or in *trans,* the latter of which resembles modification of other proteins localized to the DNA lesion. ARTD2 is also able to ADP-ribosylate the ends of the same dsDNA molecule where it is bound. This happens preferentially within a DNA molecule where the end is approximately 30 Å from the WGR binding site^29^. This indicates that the automodification of ARTD2 or PARylation of DNA may happen *in cis* and that the transferase domain is indeed mobile when the enzyme is activated.

Asn129 (ARTD1^Asn567^, ARTD3^Asn79^), located between the WGR and catalytic domains has been mapped as a key element in transferring the activation signal to the catalytic fragment^17^. We also confirmed that N129A is inactive although it retains the same nM affinity for DNA (**Fig. 4, Fig. S4, S5**). The adjacent residues, Arg153 and Asn127, also provide a link between DNA and the WGR and RD domains (**Fig. 4A**). We have shown that mutations in these residues result in loss of DNA dependent activity and specific DNA binding (**Fig. 4C**)^14^. DNA affinity of the full length protein is driven largely by the disordered N-terminus and the affinities of all the mutants generated in this study including the above also show similar single digit nM affinity for nicked phosphorylated DNA and 21-64 nM KD values for a dsDNA model (**Fig. S4**)^12,14^. An exception is Y201F, which has a slightly lower affinity also for nicked DNA (KD 21 nM). Arg153 and Asn127 interact with DNA at the 3’ side of the nick, with each other, and with a carbonyl group of Gly314 of the RD (**Fig. 4A**). Together with the Asn129 interaction, changes in these residues could result in the unfolding of some of the RD helices leading to release of the RD domain from the ART domain. Tyr201 is also important for the DNA binding of ARTD2^14^ and critical for the nicked DNA detection by ARTD2, as a Y201F mutation resulted in a 50% reduction of the catalytic activity (**Fig. 4C**).

**Figure 4.**
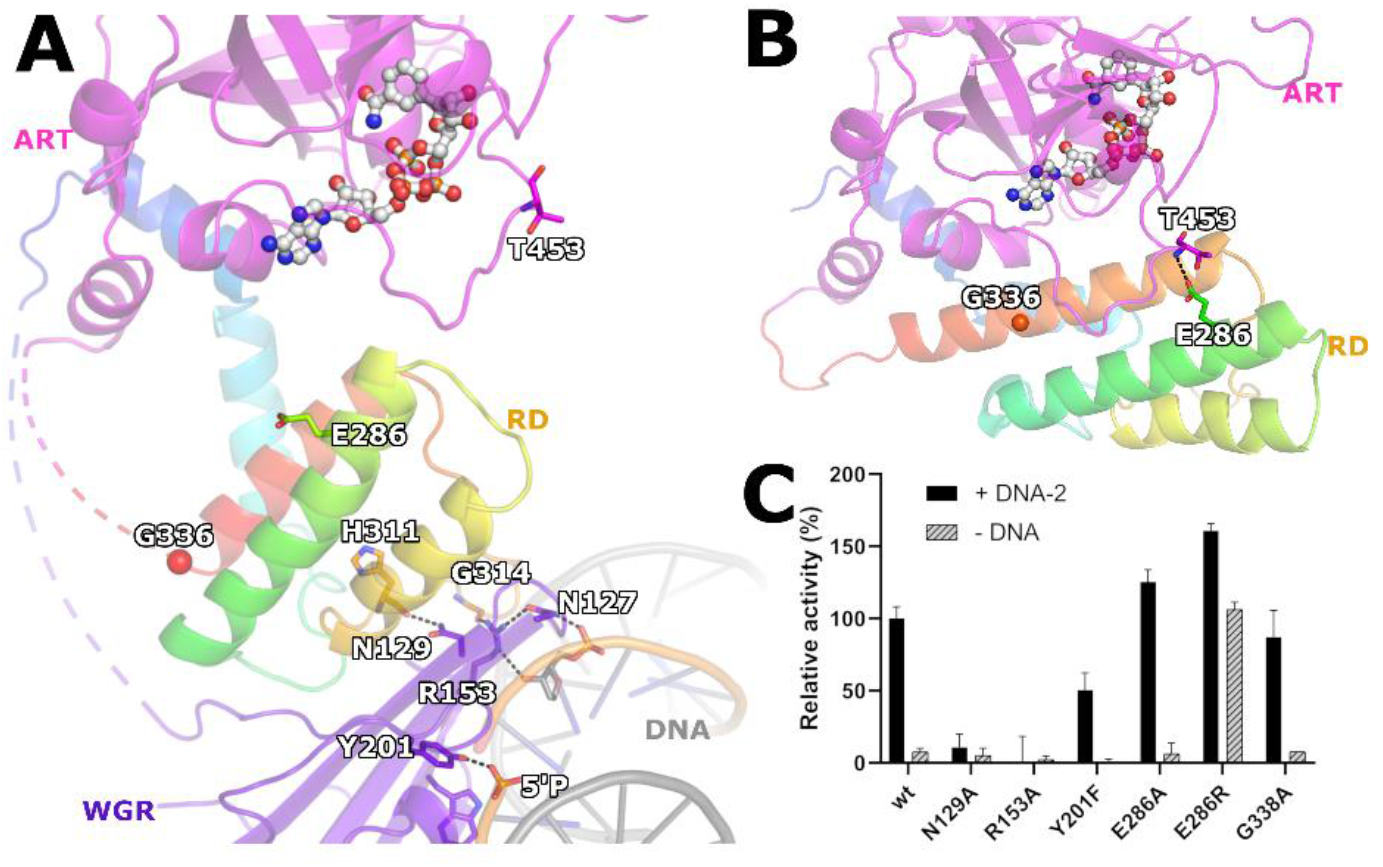
ARTD2 activation mechanism. (**A**) Upon DNA binding the interactions formed by Arg153, Asn127 and Asn129 transmit the signal for the conformational change to release the RD-ART interaction. (**B**) A model of an ARTD2 binding to DNA in an inactive conformation showing the interaction between the RD and ART domains in the inactive conformation. **(C)** DNA dependent activity assay measuring NAD^+^ consumption after incubation with ARTD2 enzymes. Data shown are means of four replicates with a SD.

To further examine the changes observed for domain interactions we designed key mutations at the RD-ART interface. We observed that the RD and ART domains have contacts at Glu286 of the RD and Thr435 backbone amides and hydroxyl located at the D-loop lining the NAD^+^ binding cleft in the inactive state (**Fig. 4B**). We rationalized that by disrupting these contacts we could create an enzyme, which would be active even in the absence of DNA. Mutant E286A only showed slightly increased activity in the presence of an activating DNA oligonucleotide, but only basal level activity in the absence of the DNA (**Fig. 4C)**. However, when reversing the charge with an incompatible E286R mutation we generated repulsion between the domains resulting in a hyperactive enzyme. The activity of the mutant is further increased by supplementing DNA indicating that the equilibrium clearly favors more the active state than the inactive state. Our effort to stabilize the helix α5 with a G338A mutation did not prevent activation (**Fig. 4C**) suggesting that a similar activation mechanism may also be possible in ARTDs which lack the glycine in the same position (ARTD1^Q717^, ARTD3^E237^).

### Distinct mode of ARTD2 in the presence of HPF1

DNA binding has been shown to be prerequisite for binding of HPF1 to the ART domain, as the RD domain of ARTD1 or ARTD2 would prevent this in the inactive conformation^27^. Based on the ART and HPF1 complex structure (PDB code, 6TX3), we modeled the position of HPF1 to the activated ARTD2. Our observed conformational changes in the ARTD2_WGR-RD-ART_ DNA complex structure indeed allow binding of HPF1 to ARTD (**Fig. 5A, Fig. S6**). This explains how, upon activation by DNA damage, the release of the ART domain allows the docking of HPF1 E284 to the catalytic core of ARTD2 modulating the specific serine ADP-ribosylation^27^. It should be noted here that in the recent cryo-EM structure the HPF1 is binding to the closed form of the ARTD2^28^ and therefore the changes enabling HPF1 binding may be more subtle (**Fig. S6**).

**Figure 5.**
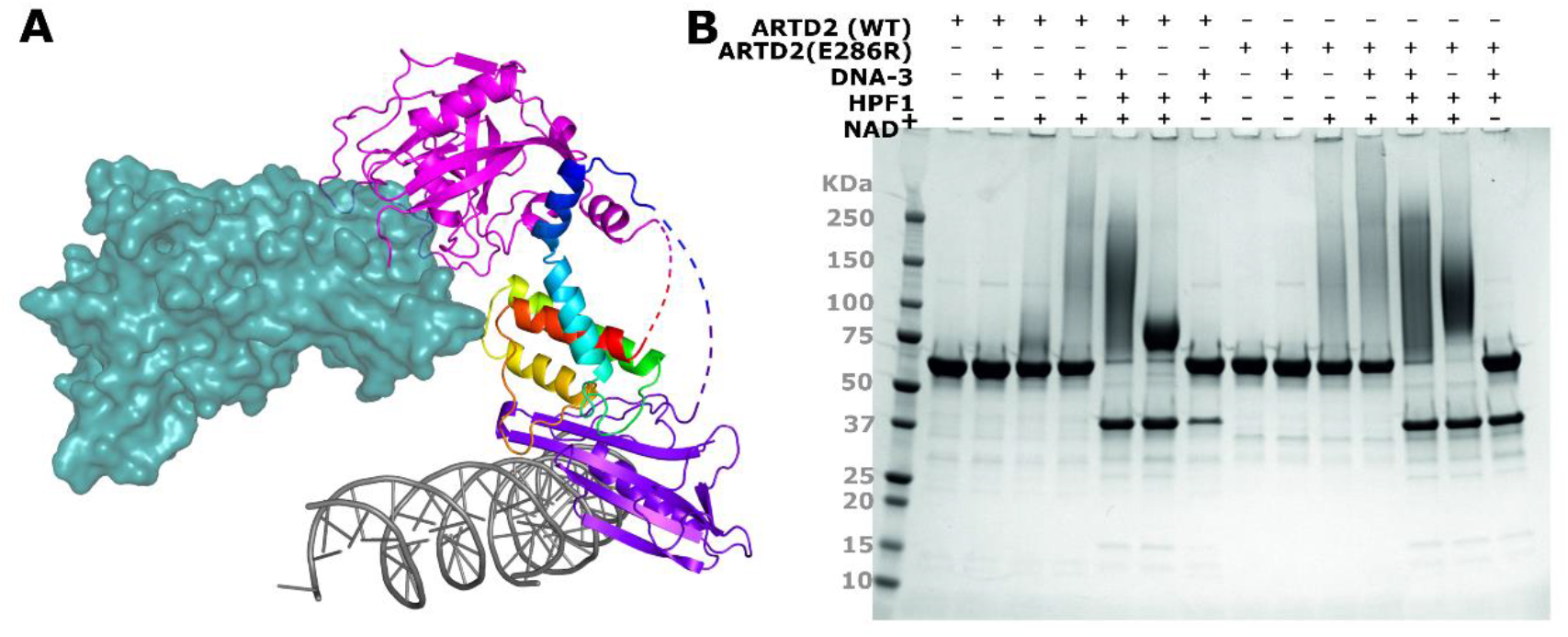
ARTD2-HPF1 complex. (**A**) Model of the ARTD2 HPF1 complex when ARTD2 is in the activated state. (**B**) Binding of HPF1 changes the nature of the resulting PAR.

We next tested the effect of ARTD2 binding in an ARTD2 automodification assay (**Fig. 5B**). In the presence of NAD^+^, ARTD2 automodification appears as a smear on an SDS-PAGE gel due to heterogeneous auto-poly-ADP-ribosylation (**Fig. 5B**). The smear is more prominent when the activating DNA-3 is present and also in the E286R mutant, both with and without DNA, in agreement with the NAD^+^ consumption assay results. The addition of HPF1 drastically affects the behavior of both WT and E286R by promoting automodification of apparently all ARTD2 enzymes present (**Fig. 5B**). The addition of HPF1 in the reaction resulted in a focused smear centered around 125 kDa. In the absence of activating DNA, HPF1 still promotes partial automodification of ARTD2 resulting in a band shift of ARTD2. Under these conditions, the WT ARTD2 automodification is more homogeneous and the protein migrates as a band centered around 75 kDa, while E286R produces species centered above 100 kDa, similar to those produced in the presence of DNA. Altogether, this suggests that HPF1 potentially contributes to the activation of ARTD2 through a mechanism involving disruption of the autoinhibitory effect of the RD domain.

## Discussion

ARTD2 is a DNA repair enzyme and its catalytic activity is highly elevated in response to cellular genotoxic stress. Here we described the structural mechanism of ARTD2 DNA damage detection and mechanism of its activation upon binding to activating DNA molecule. The current view is that the three DNA dependent ARTDs, ARTD1-3, share a similar activation mechanisms despite the distinct DNA recognition modes resulting from to the differences in their domain organization^17^. The DNA binding mode of ARTD2 has been shown to be different compared to ARTD1, which uses the zinc finger and WGR domains to coordinate its binding to DNA end while ARTD2 uses its WGR domain to create DNA end-to-end binding^14,28^. In contrast to ARTD1, the *in vitro* activity of ARTD2 is rapidly elevated only in the presence of 5’-phosphorylated DNA^12,17^.

The crystal structure described here contains an almost full length ARTD2 lacking only the disordered N-terminal responsible for nuclear localization and increased DNA affinity. The structure allows visualization of the series of events that enable ARTD2 to bind the substrate NAD^+^ upon detection of a DNA damage site. It explains previous observations of local unfolding of the RD domain, re-organization required for HPF1 binding and how ARTD enzymes can modify other proteins and themselves *in cis* and *in trans* in the context of DNA repair. A summary of the process is illustrated in **Figure 6** and it can be divided to five stages marked as i-v.

**Figure 6.**
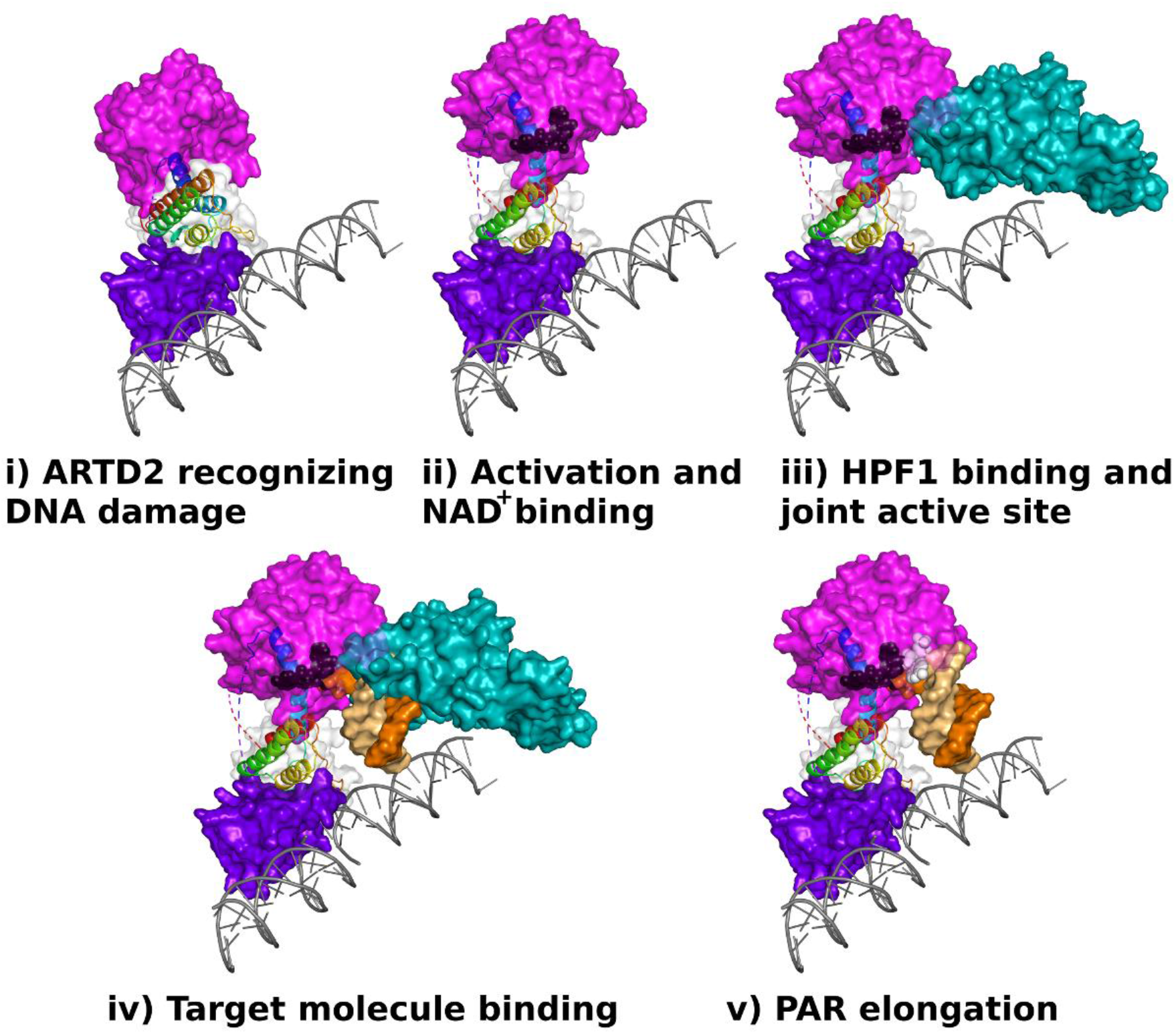
Conformational changes upon ARTD2 activation and substrate protein modification. **i)** Inactive conformation of ARTD2 when bound to DNA. **ii)** ARTD2 undergoes a conformational change in the regulatory domain α-helices allowing NAD^+^ (black) binding. **iii)** HPF1 can bind to the ARTD2 catalytic domain to form a joint active site. **iv)** Conformational changes in the regulatory domain release the catalytic domain and allow it to modify target macromolecules marked here by the second DNA molecule observed in the crystal structure. **v)** ARTD2 is able to catalyze the PAR chain elongation reaction also without HPF1, which may dissociate from the complex. Carba-NAD^+^ bound to the acceptor site is shown in white.

**i)** Initially the binding of ARTD2 to DNA is facilitated by the positively charged and disordered N-terminus and when ARTD2 recognizes a nick in the DNA, it is locked in place by bridging the DNA gap with the WGR domain^14^. The structural conformation observed for the isolated catalytic domain is very similar to the ARTD1^26^ structure in complex with DNA that was used to model inactive ARTD2. **ii)** ARTD2 binding to 5’-phosphorylated DNA induces major conformational changes in the RD domain. Previously local unfolding and exposure of these elements was demonstrated by HDXMS and by a low resolution cryo-EM structure^28^. The conformational change allows efficient binding of substrate NAD^+18^. Crystal structures of the catalytic fragment have formed a basis for structure-based drug development efforts. Recently the difference in the PARP trapping efficiency of clinical inhibitors competing with NAD^+^ binding was rationalized to result from interactions with the RD^15^. The conformational changes we observed upon DNA binding could facilitate also the development of improved drugs. **iii)** When HPF1 binds to the catalytic domain of an activated ARTD2, they form a joint active site and change the ARTD2 residue specificity from glutamate/aspartate to serine^27^. Notably, the order of ii and iii stages is not established as it was shown that NAD^+^ mimicking inhibitor was required to establish stable binding of HPF1^27^, while this was not required in the cryo-EM structure^28^. In the cryo-EM structure, the large conformational change was not observed, but in that particular case, a cluster of positively charges surface residues of HPF1 is interacting with the major groove of the nearby nucleosome DNA likely limiting the conformational flexibility (**Fig. S6**). Based on our experiments the initiation reaction in the context of automodification is very robust when HPF1 is present as all the ARTD2 proteins are modified (**Fig. 5B**). **iv)**. The large conformational changes in the regulatory domain allow binding of a substrate macromolecules that get ADP-ribosylated. In our model, a substrate protein site is marked by the second DNA molecule observed in the crystal structure, which is bound to the active site by a thymine base. Some destabilization of the HPF1 helices was reported in the low resolution cryo-EM data also for the HPF1 and this could enable substrate binding to avoid clashes with the proteins^28^. **v)** ARTD2 is able to catalyze the polymer formation alone and this happens *in vitro* (**Fig. 5B**) and it can also happen in the cell in various contexts. The so called acceptor site has been mapped based on the crystal structure of ARTD1 catalytic domain with carba-NAD^+21^. Electron density for the substrate analog is not good and subsequently it was only partially modeled in the structure. Carba-NAD^+^ is bound to a location overlapping with HPF1 binding site. This and the long polymers generated in absence of HPF1 indicate that HPF1 can dissociate at some stage from the complex and allow robust generation of long poly-ADP-ribose chains. In **figure 6**, the second DNA molecule marks the target protein position in the elongation reaction, but it is not known what the exact site of the mono-ADP-ribosylated protein is at this stage. When the polymer is elongated it likely dissociates also from the ARTD2. The sequence of these final events are not completely yet validated by experiments and substrate proteins have not been observed so far bound to the activated ARTD2 or other enzymes of the family. Our crystal structure of the open ARTD2 structure however allows visualization of these events based on current knowledge.

## Online methods

### Cloning, protein expression and purification

The cloning of the DNA constructs coding for ARTD2_FL_ isoform 1 (residues 1-583: Uniprot id. Q9UGN5) and the individual domain construct (ARTD2_WGR-RD-ART_; residues 90-583) has been previously described^12,14^. Human HPF1 was cloned into pNH-TrxT and the resulting protein after tag cleavage contains an additional S at the N-terminus. Mutagenesis of the ARTD2_FL_ enzyme and ARTD2_WGR-RD-ART_ were done using Quick change site-directed mutagenesis except for G338A, E286A and E286R mutants that were obtained by assembly PCR. Briefly, for each mutant two fragments were PCR amplified, the first comprising the 5’ end until the mutation site and a second product comprising the mutation site until the 3’ end. The two fragments were then assembled by PCR and cloned using SLIC into pNIC-Zbasic or pNH-TrxT. All clones were sequenced using the automated sequencer in the Biocenter Oulu core facility, University of Oulu, Finland. Expression and purification of proteins were done as described^12,14^. In brief, ARTD2 proteins were expressed in *E. coli* BL21 (DE3) and purifications were done in a three-step chromatography system (IMAC, Heparin and size exclusion).

For HPF1, expression was carried out in terrific broth auto-induction media (Formedium) supplemented with 0.8% glycerol (w/v) and 50 μg/mL kanamycin. The cells were grown at 37 °C with shaking until the OD600 reached 1 and then the temperature was lowered to 18°C for 16 hours. The cells were harvested by centrifugation (4000 × g, at 4 °C for 30 minutes) and suspended in lysis buffer [50mM Hepes (4-(2-hydroxyethyl)-1-piperazineethanesulfate), 500mM NaCl, 10% glycerol, 10mM imidazole, 0.5mM TCEP (tris(2- carboxyethyl)phosphine), pH 7.5] and supplemented with 0.1 mM Pefabloc (4-(2-Aminoethyl) benzenesulfonyl fluoride hydrochloride from Sigma-Aldrich). Lysis was performed by sonication and the cell debris cleared by centrifugation (30000 × g at 4°C for 30 minutes). The supernatant was filtered through a 0.45 μm filter and loaded onto a 5 ml HisTrap column (GE Healthcare). The column was washed with 180 ml of wash buffer (50mM Hepes, 500mM NaCl, 10% glycerol, 25 mM imidazole, 0.5 mM TCEP, pH 7.5). HPF1 was eluted with 50 ml of elution buffer (50mM Hepes, 500mM NaCl, 10% glycerol, 350mM imidazole, 0.5 mM TCEP, pH 7.5). The eluted sample was then diluted 1:5 in ion exchange loading buffer (25 mM Tris, 0.5 mM TCEP, pH 7.5) and loaded into a 5 ml HiTrap Q XL column (GE Healthcare) pre-equilibrated in ion exchange wash buffer (25 mM Tris, 100 mM NaCl, 0.5 mM TCEP, pH 7.5). The column was then washed with 50 ml of ion exchange wash buffer and eluted in a gradient from 100 mM to 1 M NaCl. The fusion tag was cleaved with TEV-protease (1:30 TEV:HPF1 molar ratio) at 4 °C for 16 hours. 25 mM imidazole was added to the sample and then the protein was then loaded onto a 5 mL HisTrap (GE Healthcare) and the flow through containing the cleaved proteins was collected, concentrated and further purified using Superdex 75 (30 mM Hepes, 350 mM NaCl, 10% glycerol, 0.5 mM TCEP, pH 7.5). The purified protein was pooled, concentrated, flash frozen and stored in −70 °C.

### CD

CD spectra of ARTD2_FL_ wt, N127A, N129A, E286A, E286R and G338A were recorded at 22 °C using Chiranscan CD spectroscopy (Applied Photophysics Ltd.) equipped with a temperature-regulated sample chamber. The far-UV region spectra (190-280nm) was measured in a 1 mm path length quartz cuvette. The sample concentration was 0.05-0.07 mg/mL in 10 mM sodium phosphate pH 7.4, 150 mM (NH_4_)_2_SO_4_. The data were analyzed with the Pro-Data Software suite (Applied Photophysics Ltd.).

### SDS-PAGE automodification activity assay

Automodification reactions containing 5 μg ARTD2_FL_ and equimolar concentrations of HPF1 and 5’phosphorylated hairpin oligonucleotide (DNA-3, see **Table S2**) were initiated by the addition of 1 mM NAD^+^. The reaction buffer was 50 mM Tris, 5mM MgCl_2_, pH 7.5. The samples were incubated at RT for 1 h and the reaction stopped by the addition of SDS containing buffer. The samples were then incubated 5 min at 95°C and loaded onto a Mini-Protean 4-20% TGX gel (BioRad). Following electrophoresis, the gel was stained in PageBlue protein stain solution (Thermo Scientific).

### SEC-MALS

SEC-MALS analysis was performed as described previously^14^. Briefly, 35 μM ARTD2_WGR+CAT_ and 37 μM DNA-1 were mixed in 20 mM HEPES pH 7.5, 400 mM NaCl, 0.5 mM TCEP and incubated for 1h at RT before analysis. Samples were run in an S200 increase column (GE Healthcare) and analyzed using the miniDAWN Treos II (Wyatt Technology). Mass determination was performed with ASTRA software (Wyatt Technology).

### Fluorescence activity assay

Activity assays of the ARTD2 protein were done as reported earlier^12,14,30^. 50 nM of ARTD2_FL_ wt or point mutant was mixed with 50 nM of each of the oligos (see **Table S2** for details) and 5 μM NAD^+^. Samples were incubated at RT for 15 min for the mutant comparison assay and 1 h for the DNA dependent activation assay. IC_50_ determination was performed with 40 nM ARTD2_FL_, 10 μg/ml activated DNA and 500 nM NAD^+^ and the reactions were incubated for 30 min at RT. Measurements were done in quadruplicate and repeated 3 times. Data analysis was performed with a R script using the propagate package for first order Taylor expansion uncertainty estimation and nls function to fit the Hill equation.

### Fluorescence polarization

Fluorescence polarization was performed as previously described using fluorescein tagged dumbbell DNA containing a nick and 5’-phosphate or a double-stranded DNA model with 5’- phosphate **Table S2**^12^. Measurements were done in triplicate.

### Crystallization

A 140 μL solution containing 150 μM ARTD2_WGR-RD-ART_ and 160 μM DNA-1 (purchased from Integrated DNA Technology, IDT) was incubated on ice for 30 minutes. The complex was purified with size exclusion chromatography using a Superdex^TM^ S200 Increase 10/300 GL (GE-Healthcare) column pre-equilibrated with a buffer containing 20 mM Hepes, pH 7.5, 300 mM NaCl, 0.5 mM TCEP. The fractions corresponding to the complex were collected and concentrated for crystallization. The crystallization was done using a sitting drop vapor diffusion method at +4°C and the precipitant solution was 0.1 M MES pH 6.5 and 1 M ammonium sulfate. Notably, successful crystallization required optimization of the DNA length. Our previous studies showed that the catalytic activity of ARTD2 at a physiological salt concentration of 150 mM was higher when DNAs with lengths of 10-20 base pairs (bp) were used ^12^ Based on this information, we started crystallization experiments of ARTD2_WGR-RD-CAT_ with DNAs within this range. We observed that crystallization of ARTD2WGR-RD-CAT with 12 and 15 bp DNAs produced only micro-crystals being less than 5 microns while crystallization with 16 and 20 bp DNAs produced crystals, which were large enough for data collection. However, the crystals with 20 bp DNA were very fragile and diffracted only to 8 Å in the initial data collection using the X-ray diffractometer with a rotating anode at Biocenter Oulu, Finland. However, the crystals with 16 bp DNA had much better diffraction, 4 Å, suggesting that the 16 bp dsDNA phosphorylated at one of the 5’ends (DNA-1; **Table S1**) offered a better crystal packing.

### Data collection, structure determination and refinement

In initial data collections at synchrotrons, we noticed that the diffraction quality of the crystals of the ARTD2WGR-RD-ART -DNA-1 complex was highly dependent on cryo-protectants. In addition, the crystals suffered greatly from radiation damage and the diffraction was nonuniform requiring a grid scan to locate the best diffracting positions of a single crystal. We tested PEG400 and glycerol supplemented with the mother liquor (0.1 M MES pH 6.5, 1 M ammonium sulfate) as a cryo solution, but neither of them worked well, as the crystals had a modest diffraction to 4 Å. By combining different cryo-protectants together, the diffraction quality and resolution were improved. Finally, we used 0.1 M MES pH 6.5, 1 M ammonium sulfate, 10% (v/v) glycerol, 10% (v/v) diethylene glycol, and 10% (v/v) 2-propanol as a cryosolution. However, the crystals still suffered significantly from radiation damage allowing only collection of approximately 10° oscillation with a resolution between 2.7 - 3 Å. This together with nonuniform diffraction made it impossible to obtain a complete dataset from a single crystal. In order to overcome the problem, we located the best diffracting positions of multiple crystals using grid scans and collected multiple small datasets from the positions on the beamline i24 at the Diamond Light Source (UK). The datasets were processed using XDS^31^ and their correlation in terms of unit cells and space group were analyzed using the ccCluster program^32^. Visual analysis of the dendogram in ccCluster GUI showed that the majority of the datasets were identical within a 0.2 threshold. Finally, we selected 27 datasets collected from 4 crystals for merging that yielded good statistics with a resolution of 2.8 Å in POINTLESS^33^. The data collection and structure refinement statistics are presented in **Table 1**.

Phases for the ARTD2WGR-RD-ART -DNA-1 complex structure were solved using molecular replacement with several cycles. First, the ARTD2 WGR and ART domain structures (PDB id. F6IK and 5DSY, respectively)^14,16^ together were used as search models in MRBUMP^34^ included with the protein and DNA sequences. As a result, we obtained an incomplete solution where only the WGR domain and DNA-1 were correctly placed. Next, we performed a second run using the incomplete solution and the ART domain (PDB id. 5DSY) as search models, and obtained a complete solution where the ART domain was correctly placed together with the WGR domain and DNA-1. The structure was initially refined with REFMAC5^35^ and we were able to trace the C-terminus of the WGR domain and the N-terminus of the ART domain from the structure allowing us to manually build the missing RD domain using Coot^36^. Finally, the ARTD2WGR-RD-ART DNA-1 complex structure was refined with Phenix^37^. The data collection and refinement statistics are presented in **Table 1**. The figures of the structures were made using Pymol^38^.

## Data availability

The data that support the findings of this study are available from the corresponding author upon reasonable request. Atomic coordinates and structure factors have been deposited to the Protein Data Bank under accession number 7AEO.

## Acknowledgements

We are grateful to local contacts at DLS for providing assistance in using the beamlines. This work was funded by the Jane and Aatos Erkko Foundation and the Academy of Finland (grant nos. 287063, 294085 and 319299 to LL). The use of the facilities of the Biocenter Oulu Structural Biology core facility, member of Biocenter Finland, Instruct-ERIC Centre Finland and FINStruct, as well as of “Proteomics and Protein Analysis” and Sequencing core facilities are gratefully acknowledged.

## Author information

### Contributions

LL conceived the research. EO and LL designed the research. EO and MMM, collected the X-ray diffraction data. EO solved the structure. EO, MMM, and LL refined the model. EO, MMM, AGP performed the mutagenesis and protein production. AGP performed and analyzed the binding studies. AGP performed and analyzed the activity assays. EO, LL, MMM and AGP wrote the paper.

## Ethics declarations

Competing interest

## Notes

### Competing Interest Statement

The authors have declared no competing interest.

## References

1. Schreiber, V. et al. Poly(ADP-ribose) polymerase-2 (PARP-2) is required for efficient base excision DNA repair in association with PARP-1 and XRCC1. J. Biol. Chem. 277, 23028–23036 (2002).

2. Boehler, C. et al. Poly(ADP-ribose) polymerase 3 (PARP3), a newcomer in cellular response to DNA damage and mitotic progression. Proc. Natl. Acad. Sci. U. S. A. 108, 2783–2788 (2011).

3. Miyoshi, T., Makino, T. & Moran, J. V. Poly(ADP-Ribose) Polymerase 2 Recruits Replication Protein A to Sites of LINE-1 Integration to Facilitate Retrotransposition. Mol. Cell 75, 1286–1298.e12 (2019).

4. De Vos, M., Schreiber, V. & Dantzer, F. The diverse roles and clinical relevance of PARPs in DNA damage repair: Current state of the art. Biochem. Pharmacol. 84, 137–146 (2012).

5. D’Amours, D., Desnoyers, S., D’Silva, I. & Poirier, G. G. Poly(ADP-ribosyl)ation reactions in the regulation of nuclear functions. Biochem. J. 342 (Pt 2), 249–268 (1999).

6. Audebert, M., Salles, B. & Calsou, P. Involvement of Poly(ADP-ribose) Polymerase-1 and XRCC1/DNA Ligase III in an Alternative Route for DNA Double-strand Breaks Rejoining. J. Biol. Chem. 279, 55117–55126 (2004).

7. Farrés, J. et al. Parp-2 is required to maintain hematopoiesis following sublethal γ- irradiation in mice. Blood 122, 44–54 (2013).

8. Hanzlikova, H., Gittens, W., Krejcikova, K., Zeng, Z. & Caldecott, K. W. Overlapping roles for PARP1 and PARP2 in the recruitment of endogenous XRCC1 and PNKP into oxidized chromatin. Nucleic Acids Res. 45, 2546–2557 (2017).

9. Mortusewicz, O., Amé, J.-C., Schreiber, V. & Leonhardt, H. Feedback-regulated poly(ADP-ribosyl)ation by PARP-1 is required for rapid response to DNA damage in living cells. Nucleic Acids Res. 35, 7665–7675 (2007).

10. Ahel, D. et al. Poly(ADP-ribose)-dependent regulation of DNA repair by the chromatin remodeling enzyme ALC1. Science 325, 1240–1243 (2009).

11. Gottschalk, A. J. et al. Poly(ADP-ribosyl)ation directs recruitment and activation of an ATP-dependent chromatin remodeler. Proc. Natl. Acad. Sci. U. S. A. 106, 13770–13774 (2009).

12. Obaji, E., Haikarainen, T. & Lehtiö, L. Characterization of the DNA dependent activation of human ARTD2/PARP2. Sci. Rep. 6, 34487 (2016).

13. Riccio, A. A., Cingolani, G. & Pascal, J. M. PARP-2 domain requirements for DNA damage-dependent activation and localization to sites of DNA damage. Nucleic Acids Res. 44, 1691–1702 (2016).

14. Obaji, E., Haikarainen, T. & Lehtiö, L. Structural basis for DNA break recognition by ARTD2/PARP2. Nucleic Acids Res. 46, 12154–12165 (2018).

15. Zandarashvili, L. et al. Structural basis for allosteric PARP-1 retention on DNA breaks. Science 368, eaax6367 (2020).

16. Dawicki-McKenna, J. M. et al. PARP-1 Activation Requires Local Unfolding of an Autoinhibitory Domain. Mol. Cell 60, 755–768 (2015).

17. Langelier, M.-F., Riccio, A. A. & Pascal, J. M. PARP-2 and PARP-3 are selectively activated by 5’ phosphorylated DNA breaks through an allosteric regulatory mechanism shared with PARP-1. Nucleic Acids Res. 42, 7762–7775 (2014).

18. Langelier, M.-F., Zandarashvili, L., Aguiar, P. M., Black, B. E. & Pascal, J. M. NAD+ analog reveals PARP-1 substrate-blocking mechanism and allosteric communication from catalytic center to DNA-binding domains. Nat. Commun. 9, 844 (2018).

19. Lehtiö, L. et al. Structural basis for inhibitor specificity in human poly(ADP-ribose) polymerase-3. J. Med. Chem. 52, 3108–3111 (2009).

20. Karlberg, T., Hammarström, M., Schütz, P., Svensson, L. & Schüler, H. Crystal Structure of the Catalytic Domain of Human PARP2 in Complex with PARP Inhibitor ABT-888. Biochemistry 49, 1056–8 (2010).

21. Ruf, A., de Murcia, G. & Schulz, G. E. Inhibitor and NAD+ binding to poly(ADP-ribose) polymerase as derived from crystal structures and homology modeling. Biochemistry 37, 3893–900 (1998).

22. Ruf, A., Mennissier de Murcia, J., de Murcia, G. & Schulz, G. E. Structure of the catalytic fragment of poly(AD-ribose) polymerase from chicken. Proceedigs Natl. Acad. Sci. U. S. Am. 93, 7481–5 (1996).

23. Eustermann, S. et al. Structural Basis of Detection and Signaling of DNA Single-Strand Breaks by Human PARP-1. Mol. Cell 60, 742–754 (2015).

24. Langelier, M.-F., Planck, J. L., Roy, S. & Pascal, J. M. Crystal structures of poly(ADP-ribose) polymerase-1 (PARP-1) zinc fingers bound to DNA: structural and functional insights into DNA-dependent PARP-1 activity. J. Biol. Chem. 286, 10690–10701 (2011).

25. Ali, A. A. E. et al. The zinc-finger domains of PARP1 cooperate to recognize DNA strand breaks. Nat. Struct. Mol. Biol. 19, 685–692 (2012).

26. Langelier, M.-F., Planck, J. L., Roy, S. & Pascal, J. M. Structural basis for DNA damage-dependent poly(ADP-ribosyl)ation by human PARP-1. Science 336, 728–732 (2012).

27. Suskiewicz, M. J. et al. HPF1 completes the PARP active site for DNA damage-induced ADP-ribosylation. Nature 579, 598–602 (2020).

28. Bilokapic, S., Suskiewicz, M. J., Ahel, I. & Halic, M. Bridging of DNA breaks activates PARP2-HPF1 to modify chromatin. Nature 585, 609–613 (2020).

29. Zarkovic, G. et al. Characterization of DNA ADP-ribosyltransferase activities of PARP2 and PARP3: new insights into DNA ADP-ribosylation. Nucleic Acids Res. 46, 2417–2431 (2018).

30. Narwal, M., Fallarero, A., Vuorela, P. & Lehtiö, L. Homogeneous screening assay for human tankyrase. J. Biomol. Screen. 17, 593–604 (2012).

31. Kabsch, W. XDS. Acta Crystallogr. D Biol. Crystallogr. 66, 125–132 (2010).

32. Santoni, G., Zander, U., Mueller-Dieckmann, C., Leonard, G. & Popov, A. Hierarchical clustering for multiple-crystal macromolecular crystallography experiments: the ccCluster program. J. Appl. Crystallogr. 50, 1844–1851 (2017).

33. Evans, P. Scaling and assessment of data quality. Acta Crystallogr. D Biol. Crystallogr. 62, 72–82 (2006).

34. Keegan, R. M. & Winn, M. D. MrBUMP: an automated pipeline for molecular replacement. Acta Crystallogr. D Biol. Crystallogr. 64, 119–124 (2008).

35. Murshudov, G. N. et al. REFMAC5 for the refinement of macromolecular crystal structures. Acta Cryst D 67, 355–367 (2011).

36. Emsley, P., Lohkamp, B., Scott, W. G. & Cowtan, K. Features and development of Coot. Acta Crystallogr. D Biol. Crystallogr. 66, 486–501 (2010).

37. Afonine, P. V. et al. Towards automated crystallographic structure refinement with phenix.refine. Acta Crystallogr. D Biol. Crystallogr. 68, 352–367 (2012).

38. The PyMOL Molecular Graphic System. (Schrödinger, LLc).

