## Supplementary information for "Activation of ARTD2/PARP2 by DNA damage induces conformational changes relieving enzyme autoinhibition"

**Affiliations:**
[1]Faculty of Biochemistry and Molecular Medicine & Biocenter Oulu, University of Oulu, Oulu, Finland.

**Content:**

**Table S1**. Oligonucleotides used in the study.
**Fig. S1.** SEC-MALS of ARTD2$_{WGR\text{-}RD\text{-}ART}$ DNA-1 complex.
**Fig. S2.** ARTD2 inhibition by thymine derivatives and activation by DNA models.
**Fig. S3.** ARTD1 with substrate analog BAD.
**Fig. S4.** ARTD2 affinity to DNA models.
**Fig. S5.** CD spectra of ARTD2$_{FL}$ wt and mutants.
**Fig. S6.** Inactivated ARTD2 model is not compatible with HPF1 binding.

**Table S1. Oligonucleotide sequences used in this study.**

| Schematic | Oligonucleotides | Sequence |
|---|---|---|
| 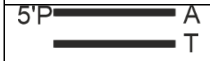  | DNA-1 | Forward: 5’phosphate GAC GAC CCG GAG CAC A 3’<br>Reverse: 3’ TGT GCT CCG GGT CGT C 3’ |
| 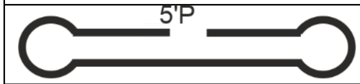  | DNA-2 | 5'phosphate GGG TCT TTT GAC CCT CGA GCT TTT GCT CGA 3’ |
| 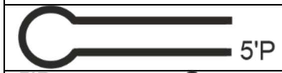  | DNA-3 | 5'phosphate GCC TAG CTA CGT AGC TAG GC 3’ |
| 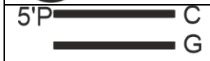  | DNA-4 | Forward: 5’phosphate GAC GAC CCG GAG CAC C 3’<br>Reverse: 5’ GGT GCT CCG GGT CGT C 3’ |
| 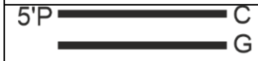  | DNA-5 | Forward: 5’phosphate GAC GCA ACC CGG AGA ACA CC 3’<br>Reverse: 5’GGT GTT CTC CGG TTG CGT C 3’ |
| 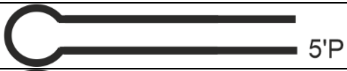  | DNA-6 | 5’phosphate GCC TAG CTA CGT AGC TAG GC 3’ |
| 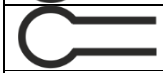 | DNA-7 | 5’ GCC TAT ATA GGC 3’ |
| 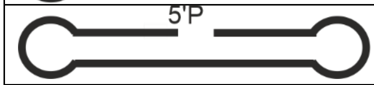  | DNA-8 | 5’phosphate GGA AGT CTT T(Fluorescein) TGA CTT CCT CGA AGC TTT TGC TTC GA 3’ |
| 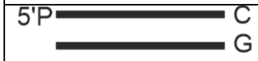  | DNA-9 | 5´phosphate GAC GCA ACC CGG AGA ACA CA 3’<br>Reverse: 5’ TGT GTT CT(Fluorescein)C CGG GTT GCG TC 3’ |

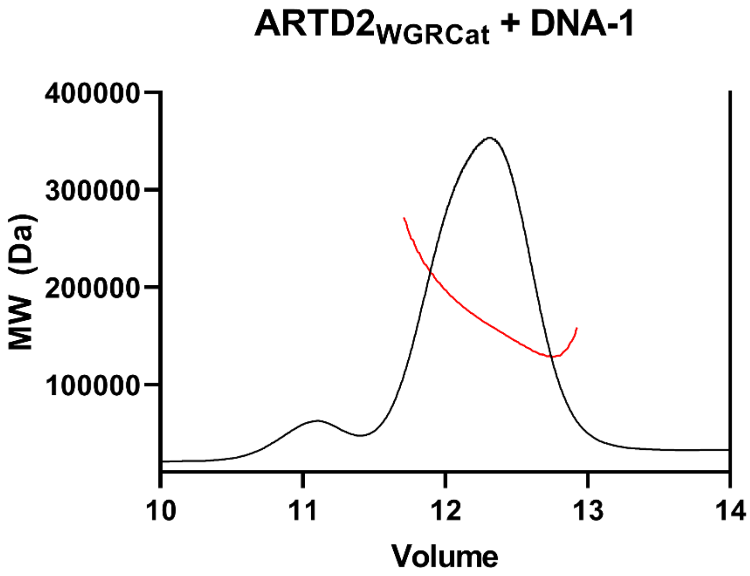


**Fig. S1. SEC-MALS of ARTD2$_{\mathrm{WGR\text{-}RD\text{-}ART}}$ DNA-1 complex.** 35 µM of ARTD2$_{\mathrm{WGR\text{-}RD\text{-}ART}}$ was mixed with 37 µM of DNA-1 and the complex was analyzed using SEC-MALS. The experimental MW determined at 12.6-12.8 ml elution volume was 130 kDa against a theoretical molecular weight of the 2:2 complex of 133 kDa.

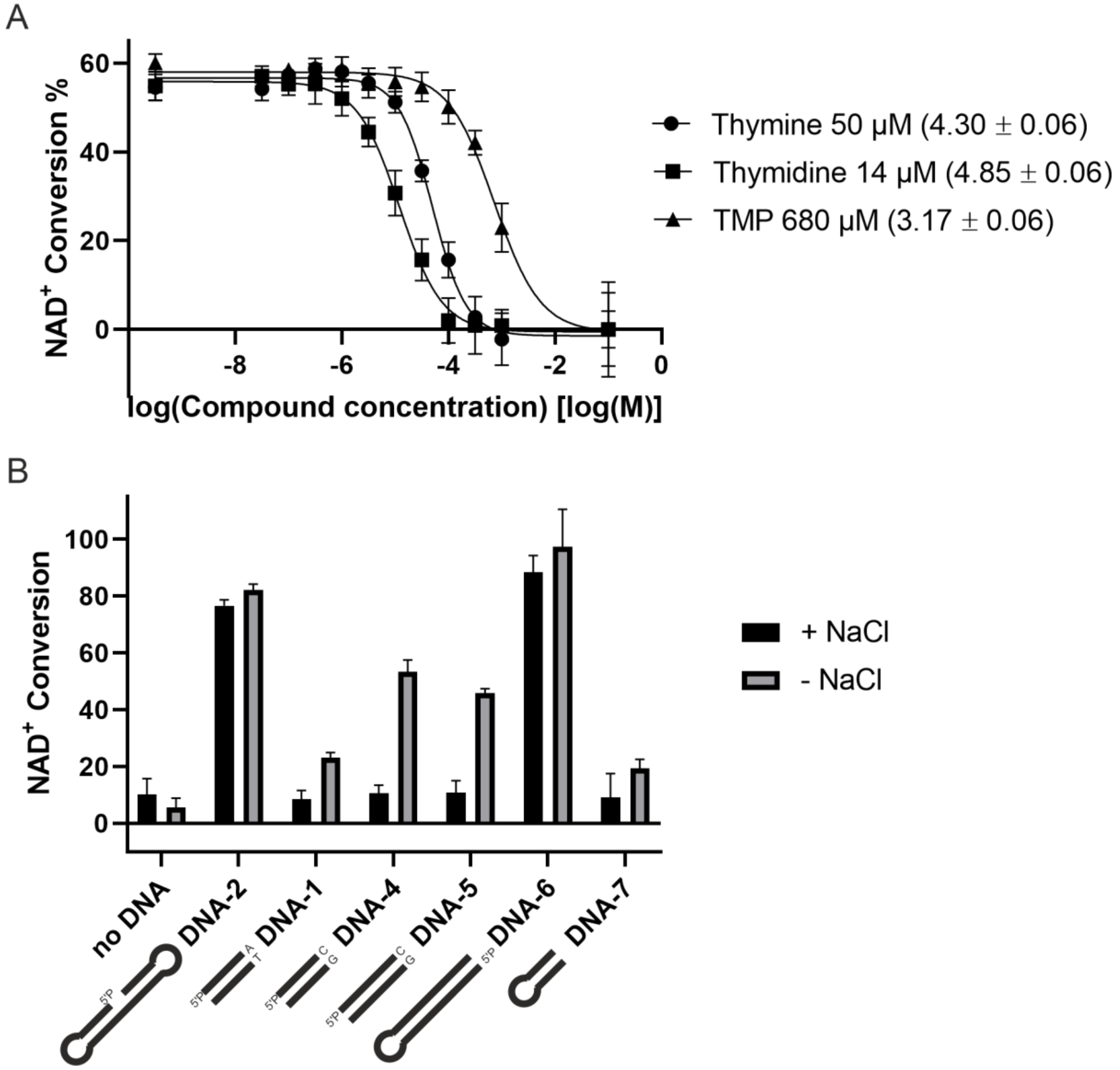


**Fig. S2.** ARTD2 activation by DNA. (**A**) $IC_{50}$ determination of Thymine, Thymidine and TMP with $ARTD2_{FL}$. Example curves with data in means and standard deviations are shown. Calculated values correspond to $IC_{50}$ ($pIC_{50}$ ± S.D.) (**B**) Catalytic activity of ARTD2 in the presence of the oligonucleotide used in crystallization (DNA-1) is very low. Replacement of the terminal A-T pair on DNA-1 with C-G (DNA-4) as well as longer double-stranded DNA terminated in C-G (DNA-5) promote activation of ARTD2 in the absence of NaCl. 5´-phosphorylated dumbbell and hairpin DNA (DNA-2 and DNA-6) used as positive controls and non-phosphorylated DNA-7 used as negative control.  Data shown are means and standard deviations of quadruplicates.

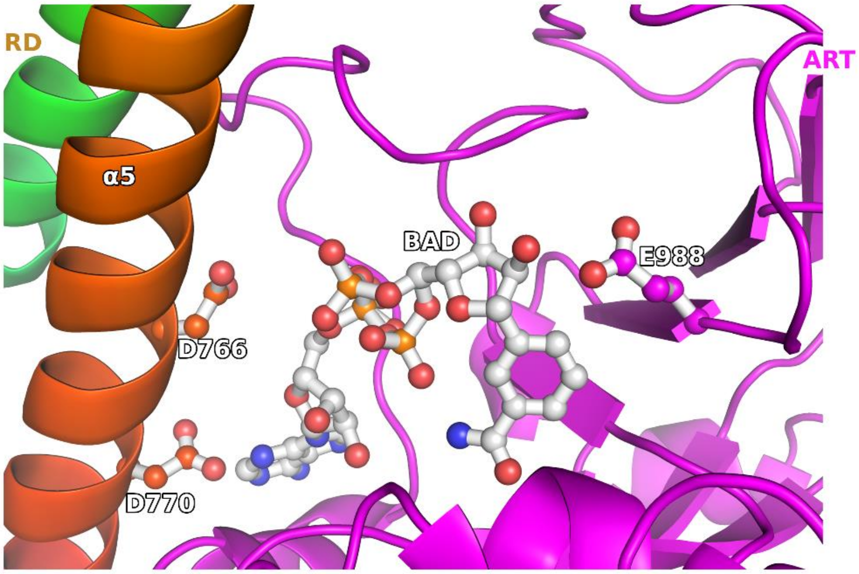


**Fig. S3.** Substrate analog BAD from the transferase domain structure (PDB id. 6BHV) superimposed to the ARTD1-DNA complex structure (PDB id. 4DQY) shows the steric hindrance between BAD and the RD domain especially with Asp770 of helix α5 (**Fig. 3**). Catalytic E988 and the negatively charged D766 located near the BAD phosphate groups are shown in sticks.

**A**

| | WT | N129A | R153A | Y201F | E286A | E286R | G338A |
|---|---|---|---|---|---|---|---|
| DNA-8 | 7.8 | 6.9 | 8.2 | 20.8 | 6.9 | 6.7 | 7.5 |
| DNA-9 | 21 | 45 | 64 | 47 | 26 | 25 | 30 |

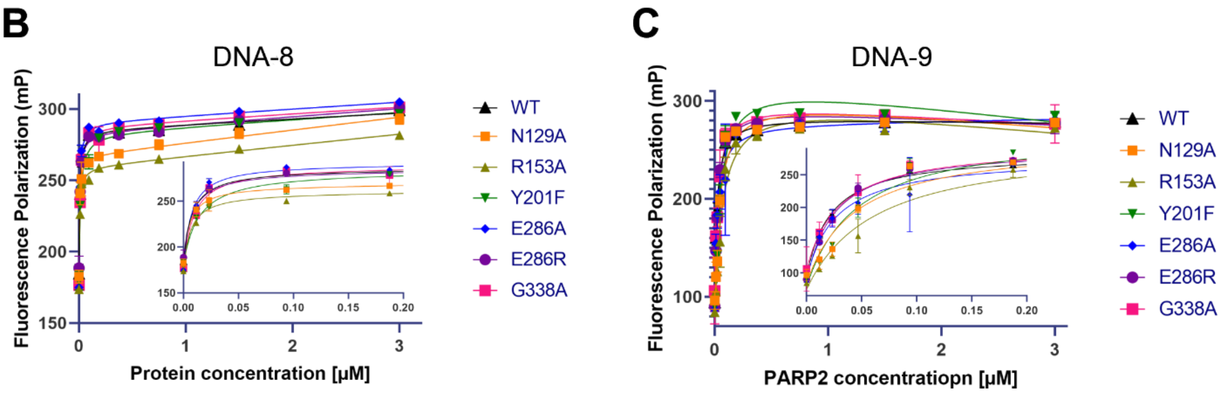


**Fig. S4.** Fluorescence polarization assay to determine the affinity of $ARTD2_{FL}$ mutants to DNA. (**A**) Summary table with estimated $K_D$ in nM. (**B**) Fluorescence polarization of dumbbell nicked and 5’-phosphorylated DNA (DNA-8) with different $ARTD2_{FL}$ mutants (**C**) fluorescence polarization of 5’-phosphorylated double-stranded blunt end DNA (DNA-9) with $ARTD2_{FL}$ mutants. The low concentration range is shown as insets in the B and C panels. Example curves are shown in panels B and C with data points as means and standard deviations of triplicates.

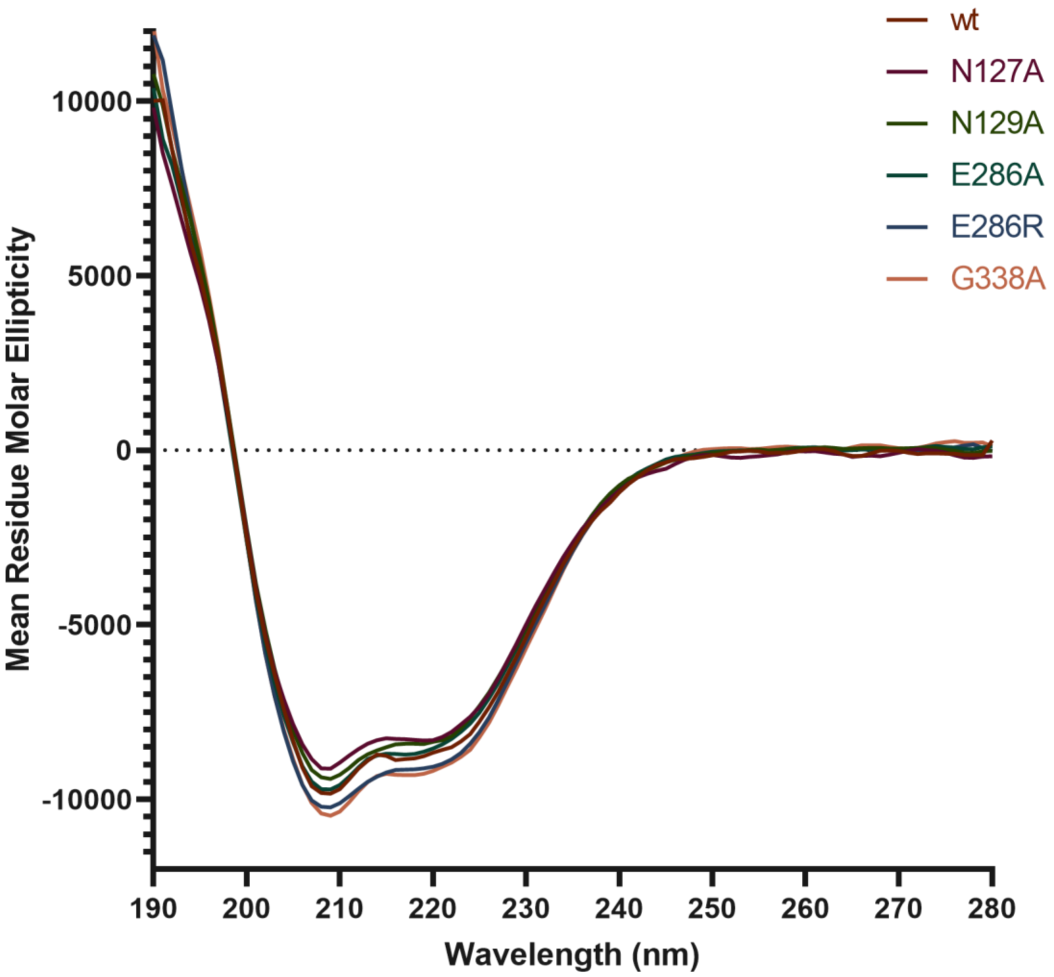


**Fig. S5**. CD spectra of $ARTD2_{FL}$ wt and mutants.

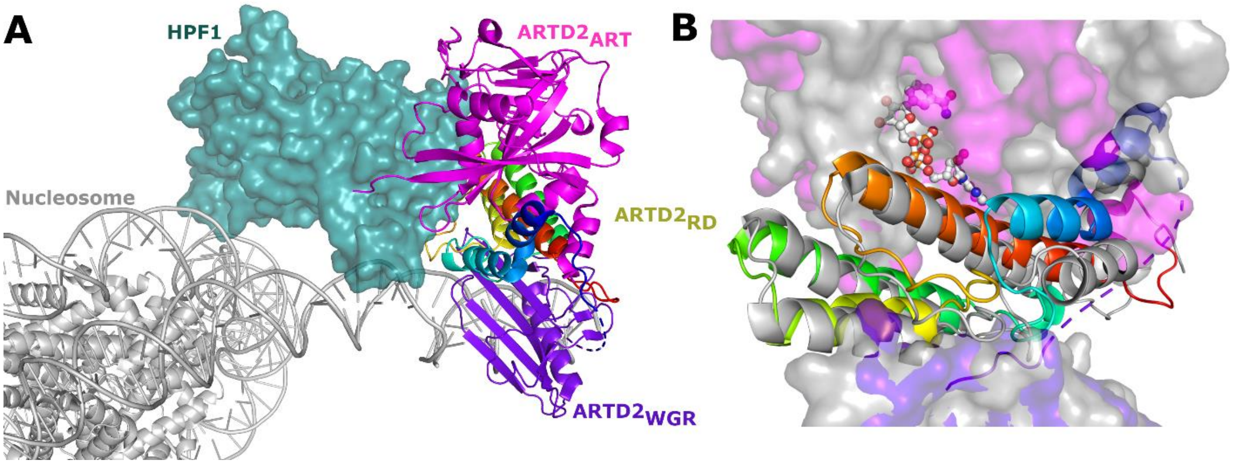


**Fig. S6**. ARTD2-HPF1 complex. (**A**) Cryo-EM structure of ARTD2-HPF1-nucleosome complex (PDB id.6X0N). (**B**) A model of an ARTD2 binding to DNA in an inactive conformation. The model was generated based on an ARTD1 crystal structure (PDB id. 4DQY) and by superimposing a ARTD2 catalytic fragment crystal structure (PDB id. 4TVJ) and a ARTD1 substrate analog BAD structure (PDB id. 6BHV). This model was superimposed with the Cryo-EM structure of ARTD2 (PDB id. 6X0N, grey).
